# Cannabinoids regulate an insula circuit controlling water intake

**DOI:** 10.1101/2022.03.18.484736

**Authors:** Zhe Zhao, Ana Covelo, Arojit Mitra, Marjorie Varilh, Yifan Wu, Débora Jacky, Astrid Cannich, Luigi Bellocchio, Giovanni Marsicano, Anna Beyeler

## Abstract

The insular cortex, or insula, is a large brain region involved in the detection of thirst and the control of water intake. However our understanding of the topographical, circuit and molecular mechanisms the controlling water intake within the insula remains parcellated. We found that type-1 cannabinoid receptors (CB_1_) within the insular cortex participate to the regulation of water intake, and deconstructed circuit mechanisms of this control. Topographically, we revealed that the activity of excitatory neurons in both anterior (aIC) and posterior (pIC) insula increases in response to water intake, yet removal of CB_1_ receptors only in the pIC decreases water intake. Interestingly, we found that CB_1_ receptors are highly expressed in insula projections to the basolateral amygdala (BLA), while undetectable in the neighboring central part of the amygdala. Thus, we imaged the neurons of the anterior or posterior insula targeting the BLA (aIC-BLA and pIC-BLA), and found they oppositely respond to water intake, respectively decreasing and increasing their activity upon water drinking. Consistently, chemogenetic activation of pIC-BLA neurons decreased water intake. Finally, we uncovered CB_1_-dependent short term synaptic plasticity (depolarization-induced suppression of excitation, DSE) selectively in pIC-BLA, compared to aIC-BLA synapses. Altogether, our results support a model where CB_1_ signaling in the pIC-BLA pathway exerts a positive control on water intake.

## INTRODUCTION

The brain perceives internal thirst states and subsequently drives drinking behavior to maintain bodily fluid homeostasis [1–6]. Imaging studies in humans and rodents suggest that the insular cortex (IC) responds to thirst-linked states by increasing its global neural activity [7–9]. However, IC neurons display heterogeneous responses upon thirst, suggesting that subpopulations of IC neurons differently encode body liquid states [10, 11]. Divergent coding of thirst states within the IC has been supported by the observation that optogenetic activation of IC projections to different amygdala nuclei lead to facilitation or suppression of licking behavior [12, 13], although optogenetic activation or inhibition of all excitatory neurons of the anterior insular cortex did not affect drinking behavior [14].

Possibly by regulating presynaptic activity, type-1 cannabinoid receptors (CB_1_) participate in the control of water intake [15–20]. In this study, we show that CB_1_ receptors expressed in IC neurons selectively modulate output pathways, which we found to differentially control water intake. Using i*n vivo* calcium imaging, we revealed that the activity of excitatory neurons in both anterior (aIC) and posterior (pIC) insula increases in response to water intake, yet removal of CB_1_ receptors expressed only in the pIC decreases water intake. Interestingly, CB_1_ receptors are highly expressed in insula projections to the basolateral amygdala (BLA), and *in vivo* calcium imaging revealed that the activity of aIC and pIC neurons targeting the BLA (aIC-BLA and pIC-BLA) respectively decreased and increased upon water drinking. Consistently, chemogenetic activation of pIC-BLA neurons decreased water intake after thirst induction. Finally, *ex vivo* whole-cell patch-clamp recordings uncovered CB_1_-dependent depolarization-induced suppression of excitation (DSE) selectively in pIC-BLA, compared to aIC-BLA synapses. Altogether, our data support a model where CB_1_ signaling in the pIC-BLA pathway exerts a negative control on water intake.

## RESULTS

### Neural activity in both anterior and posterior insular cortices (IC) increases in response to water drinking

To monitor the responses of topographically distinct neuronal populations of the IC to water intake, we used fiber photometry in freely moving mice in combination with precise recording of water licking behavior. We analyzed calcium signals in excitatory neurons of the anterior or posterior IC (aIC or pIC, respectively) of mice previously locally injected with an adeno-associated virus (AAV) carrying the gene coding for the calcium indicator GCaMP6f under the control of the CaMKII promoter (See Methods and **Figures 1A, S1A, and S1B**). Mice could freely access water in the center of an open field apparatus while calcium signal and licking behavior were recorded (**Figures 1B and 1C**). Experiments were conducted under dim red light conditions, decreasing anxiety-related effect of the open field. Remarkably, the activity of glutamatergic neurons of both aIC and pIC increased during licking behavior (**Figures 1C-1G**). Interestingly, the calcium signals at the end of each drinking bout (late licks phase) did not change after drinking (postlicking phase, **Figures S1C-1F**), suggesting that after drinking the activity of aIC and pIC glutamatergic neurons remains higher than before water consumption.

**Figure 1.**
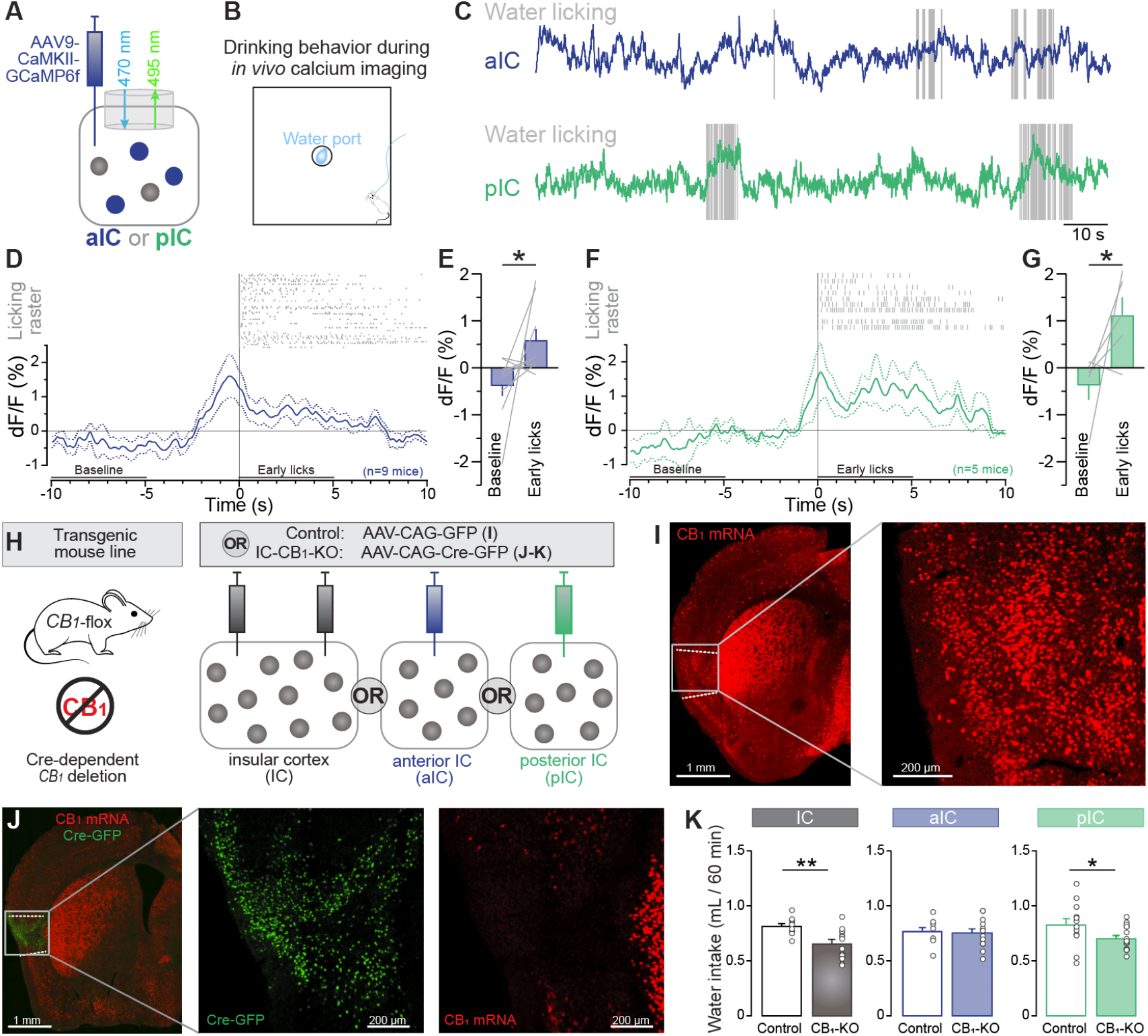
Neural activity of aIC and pIC excitatory neurons increases during drinking behavior and CB_1_ receptors in pIC cells controls water intake. **(A)** Diagram of the stereotaxic surgery for viral expression of the calcium indicator GCaMP6f in aIC or pIC excitatory neurons, and for *in vivo* fiber photometry. **(B)** Diagram of *in vivo* calcium imaging during water consumption. **(C)** Representative GCaMP6f signal recorded in the aIC (top, blue trace) or pIC (bottom, green trace), overlaid on water licking behavior (grey vertical lines). **(D)** Lick raster during drinking bouts above the averaged calcium signal of aIC glutamatergic neurons during those drinking bouts and aligned to the first lick of each bout. The signal from −10 to −5 s and from 0 to 5 s is considered as baseline and ‘early lick’ signals respectively. dF/F represents the fluorescent changes from the mean level of the entire recording time series. Dashed lines represent mean ± SEM. **(E)** The average calcium signal in aIC glutamatergic neurons is higher during ‘early licks’ compared to baseline, (two-tailed paired *t*-test, n=9, *p=0.0152). **(F)** Raster of water licking behaviors above the average calcium signal in glutamatergic pIC neurons, aligned to the first lick of the bout. **(G)** Average neural activity during ‘early licks’ in pIC glutamatergic neurons is higher than during baseline (two-tailed paired *t*-test, n=5, *p=0.0212). **(H)** Experimental strategy to knock out CB_1_ receptors in the cells of the entire IC, the aIC, or the pIC. **(I)** CB_1_ mRNAs in insular neurons of a control mouse (injected with the control virus). **(J)** Knock out of CB_1_ mRNAs in insular neurons after expressing the Cre recombinase. Note the absence of red CB_1_ mRNA signal (red) in presence of the Cre-GFP (green). **(K)** Drinking behavior induced by hypertonic NaCl i.p. injection is decreased after *CB_1_*-KO in the entire IC (n=11 control, n=13 *CB_1_-KO,* two-tailed unpaired *t*-test, **p=0.001) and *CB_1_*-KO in the pIC (n=13 control, n=16 *CB_1_*-KO, two-tailed unpaired *t*-test, **p=0.04). Drinking remained unchanged after *CB_1_*-KO in the aIC. Data in **(E, G, and K)** are shown as the mean ± SEM. For histological verifications, see **Figure S1**.

### CB_1_ receptors in posterior insula neurons regulate water intake

CB_1_ receptors have been involved in the control of water consumption [15–20], and systemic genetic removal or pharmacological blockade of CB_1_ receptors decreases water intake [15, 16]. As we observed increased activity in IC excitatory neurons when mice are drinking water, we hypothesized that selective deletion of CB_1_ receptors in the entire IC, the aIC or the pIC would differentially alter water consumption. To achieve conditional deletion of CB_1_ receptors in specific regions of the IC, an AAV coding for the Cre recombinase or the fluorescent reporter EGFP under the synthetic CAG promoter (AAV-CAG-Cre or AAV-CAG-EGFP), was injected into the entire IC, the aIC, or the pIC of *CB_1_*-Flox mice (**Figures 1H, S1G-S1I**) as reported previously [21, 22]. Fluorescent CB_1_ mRNAs *in situ* hybridization (FISH) [16, 23–25] revealed that the transcript was strongly expressed in the IC of control mice (**Figure 1I**), and absent in mice expressing Cre in the IC (**Figure 1J**). Removal of CB_1_ receptors from the entire IC or from the pIC, but not from the aIC, decreased water intake induced by a systemic hypertonic administration of NaCl (**Figure 1K**). Thus, CB_1_ receptors in the pIC appear to play a specific role in the control of water intake.

### CB_1_ receptors of insular neurons are specifically present at basolateral amygdala (BLA) terminals

Since CB_1_ receptors are mainly located at the presynaptic terminals [26–29] and IC sends intense projections to BLA and CeA [12, 13, 30–32], we next examined the distribution of insula CB_1_ receptors on axonal terminals in the amygdala including the BLA and CeA. To observe CB_1_ receptors of IC neurons specifically, we took advantage of a rescue approach to selectively re-express CB_1_ receptors in IC cells in Stop-*CB_1_* mutant mice, where CB_1_ receptor expression is prevented by a “floxed-stop” cassette, except in the presence of the Cre recombinase [16, 33, 34] (**Figure 2A**). CB_1_ receptors were re-expressed in both aIC and pIC in the mutant mice through injections of the AAV-CAG-Cre virus into IC (IC-*CB_1_*-Rescue, **Figures 2B and 2C**), but not in the control mice (**Figures S2A-S2F**). We then quantified CB_1_ receptors in IC axonal terminals within the amygdala and found that the receptors are abundantly present in the BLA and virtually absent in the CeA of IC-*CB_1_*-Rescue mice (**Figures 2D and 2E**). Importantly, there was no expression of CB_1_ receptors in the control mice (**Figures S2B-S2F**). These data reveal that CB_1_ receptors on axonal terminals of IC neurons are highly expressed in the BLA, but not in the CeA, although IC neurons intensively project to both BLA and CeA [12, 13, 30–32].

**Figure 2.**
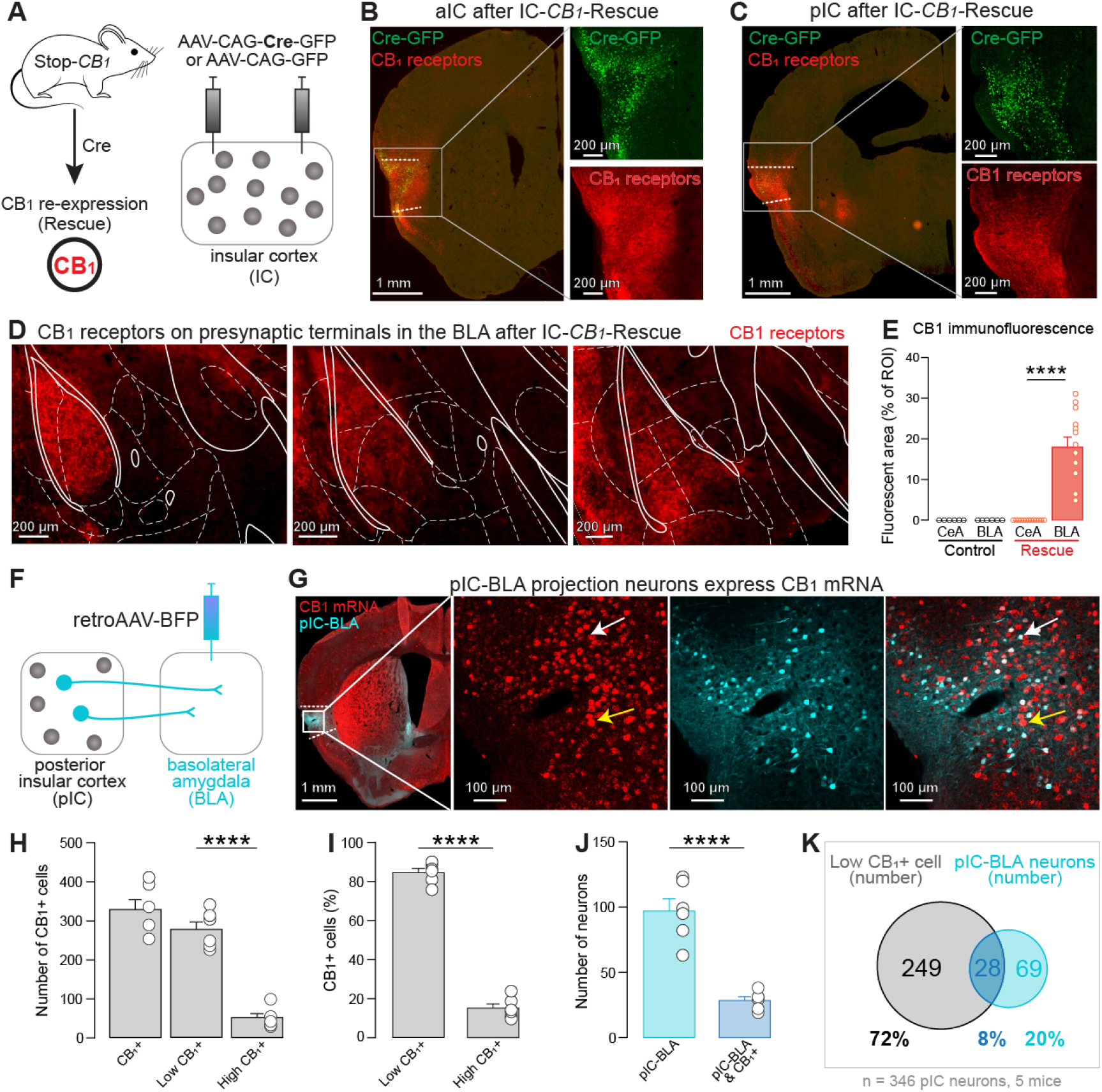
CB_1_ receptors of insula neurons are located on presynaptic terminals in the basolateral amygdala. **(A)** CB_1_-rescue experimental design to express CB_1_ receptors selectively in IC cells of Stop-*CB_1_* mice. **(B-C)** Expression of CB_1_ receptors in the aIC **(B)** and pIC **(C)** after Cre recombinase expression. **(D**-**E)** CB_1_ receptors of IC neurons are presynaptically present in the BLA but not the CeA. n=6 CeA-Control, n=6 BLA-Control, n=13 CeA-Rescue, n=13 BLA-Rescue, one-way ANOVA, F(3,34)=32.98, ****p<0.0001; CeA-Rescue vs. BLA-Rescue Tukey’s post-hoc test ****p<0.0001. **(F)** Diagram of the approach to identify IC-BLA projection neurons by injecting retroAAV-BFP into the BLA. **(G)** CB_1_ mRNAs (red) in IC-BLA projection neurons (cyan) revealed by CB_1_ fluorescent in situ hybridization (FISH) coupled with BFP immunostaining. White and yellow arrows respectively indicate cells with low and high intensity of red fluorescence which are named low-CB_1_ and high-CB_1_ positive cells. **(H)** Quantification of CB_1_-positive cell number in the pIC. High-CB_1_ cells are largely more numerous than low-CB_1_ cells in the pIC. n=6 in each group, one-way ANOVA, F(2,15)=57.09, ****p<0.0001; low**-**CB_1_ vs. high-CB_1_ cell number in pIC Tukey’s post-hoc test ****p<0.0001. **(I)** High-CB_1_ and low-CB_1_ cells represent 15% and 85% of the CB_1_-positive neurons, respectively. n=6 in each group, two-tailed paired *t*-test, ****p<0.0001. **(J)** Quantification of pIC-BLA projection neurons, along with CB_1_-positive pIC-BLA projection neurons. n=6 in each group, two-tailed paired *t*-test, ****p<0.0001. **(K)** Venn diagram of low-CB_1_ positive cell number (grey) and IC-BLA projection cell number (cyan). The overlap of grey and cyan circles represents the number of CB_1_-positive IC-BLA projection neurons. The average of the amount of low CB_1_+ cells plus pIC-BLA projection neurons per section is 346. The percentage of CB_1_-positive cells colocalized with IC-BLA projection neurons among these cells in both aIC and pIC is 8%. Data in **(E)** and **(H)** are shown as the mean ± SEM, and were analyzed by one-way ANOVA test. For more relevant information, see **Figure S2** and **Table S1**.

As we found that CB_1_ receptors in pIC cells regulate drinking behavior and insula CB_1_ receptors are selectively located on insula axon terminals in the BLA, we then quantified the number of pIC-BLA projection neurons expressing CB_1_ mRNAs. To identify IC-BLA projection neurons, a retrograde AAV coding for the blue fluorescent protein (BFP) was injected into the BLA. After viral expression, double staining of CB_1_ mRNA by FISH and of BFP by immunostaining was performed [16, 23–25] (**Figure 2F**). Here, BFP signals are shown with pseudo cyan and were found to selectively colocalize with CB_1_ mRNAs in the pIC (**Figure 2G**). As high intensity CB_1_-positive cells (high-CB_1_) are putative inhibitory interneurons and low intensity CB_1_-positive cells (low-CB_1_) can be either inhibitory or excitatory neurons [25], we quantified high and low intensity CB_1_-positive cells in the pIC from 5 mice. We found an average of 328 CB_1_-positive cells in the pIC per section (**Figure 2H**). The average number of high and low CB_1_ cells were 51 and 277 in the pIC per section (15% and 85%, **Figures 2H and 2I**). We then computed the number of pIC-BLA projection neurons in pIC sections and found an average of 97 pIC-BLA neurons per section. Low-CB_1_ cells colocalized with retrograde staining in 28 pIC-BLA neurons per section (**Figures 2J and 2K**). The fraction of low-CB_1_-pIC-BLA neurons represents 10% of all low-CB_1_ cells, 29% of all pIC-BLA neurons, and 8% of all low-CB_1_ cells and all pIC-BLA projection neurons together (**Figure 2K**). These results reveal that the pIC contains CB_1_-positive cells, IC-BLA projection neurons, as well as low-CB_1_-IC-BLA neurons.

### Insula neurons targeting the BLA respond to water drinking behavior

The BLA is one of the main targets of IC projection neurons [13, 30, 31], although their role in the control of water intake remains unclear. As we found that pIC excitatory neurons are active during water licking, that CB_1_ receptors in pIC cells regulate drinking behavior and that CB_1_ mRNA is expressed in pIC-BLA projection neurons, we hypothesized that pIC-BLA projection neurons selectively respond to water consumption and control water intake. To address this question, we first investigated how IC-BLA projections neurons respond to drinking behavior. We applied a double viral approach injecting a retrograde virus carrying the gene coding for the Cre recombinase (CAV2-Cre) into BLA and a Cre-dependent AAV including the gene coding for the calcium indicator GCaMP6m, in the aIC or pIC [12, 30, 35] (**Figure 3A**).

**Figure 3.**
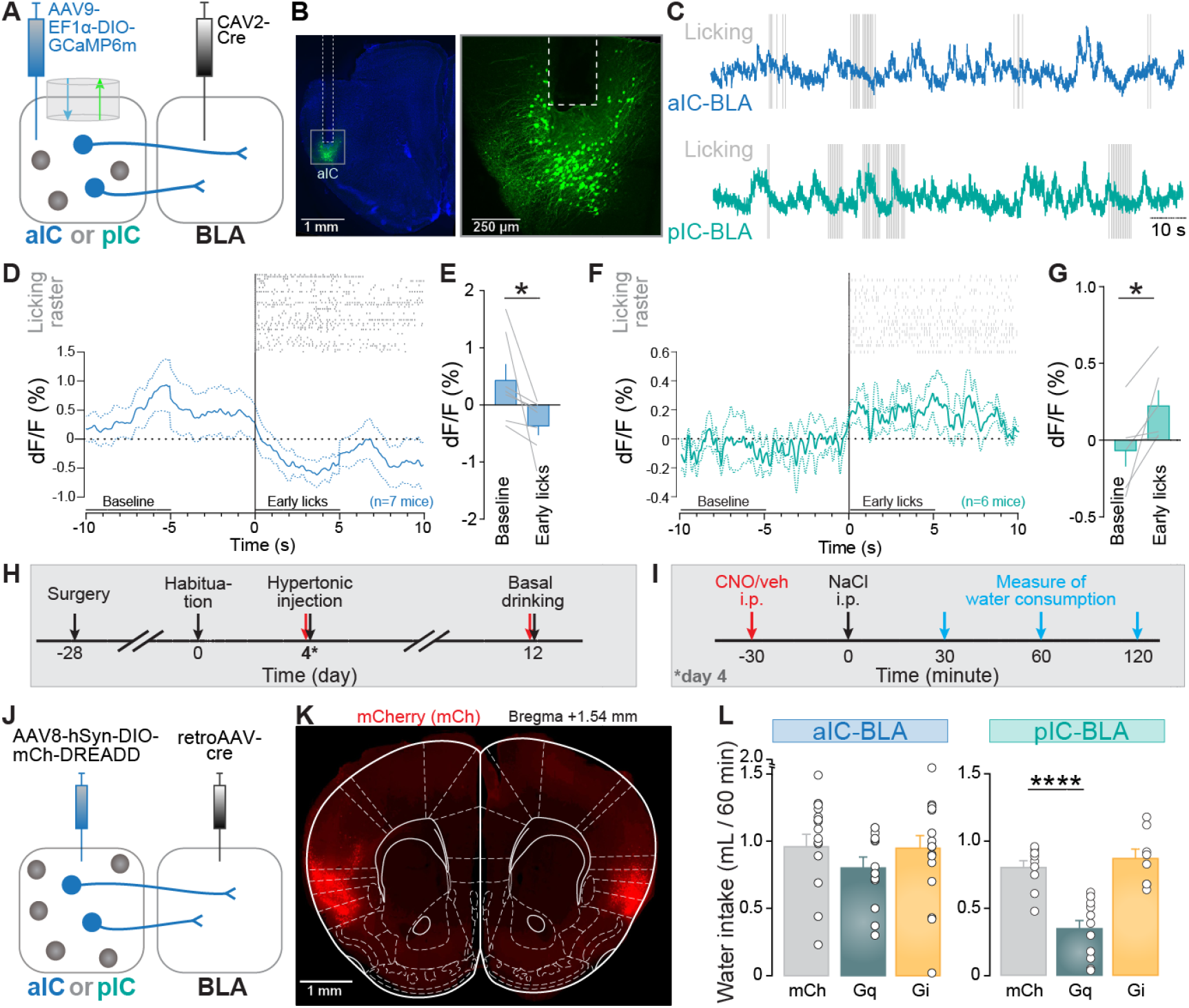
aIC-BLA and pIC-BLA neurons differentially respond to water consumption and pIC-BLA activation decreases water intake. **(A)** Combinatorial viral approach to express GCaMP6m in aIC-BLA or pIC-BLA projection neurons and fiber implantation aIC or pIC. **(B)** Traces of GCaMP6m signal of aIC-BLA projection neurons recorded in the aIC (blue trace) or pIC-BLA projection neurons recorded in the pIC (turquoise trace) overlaid on water licking events (grey vertical lines). **(C)** Confocal images of aIC-BLA projection neurons expressing GCaMP6m and track of the optical fiber implant at the aIC. **(D)** Licking raster during drinking bouts above the average calcium signal of aIC-BLA neurons. Each bout is aligned to the first lick, and the signals from −10 to −5 s and from 0 to 5 s are considered as baseline and ‘early lick’ signals respectively. dF/F represents the fluorescent changes from the mean level of the entire recording time series. Dashed lines are mean ± SEM. **(E)** The average calcium signal of aIC-BLA projection neurons is lower during early licks compared to baseline (two-tailed paired *t*-test, n=7 mice, *p=0.0156). **(F)** Licking raster above calcium signal of pIC-BLA projection neurons. **(G)** The signal is higher during the ‘early licks’ period compared to baseline (two-tailed paired *t*-test, n=6 mice *p=0.0459). **(H)** Experimental timeline of chemogenetic manipulations of aIC-BLA and pIC-BLA neurons. **(I)** Protocol to test the role of aIC-BLA and pIC-BLA neurons on NaCl-induced water intake. **(J)** Dual viral approach to express Gq or Gi DREADDs fused to mCherry (hM4Dq-mCh or hM3Di-mCh), or mCherry alone, in aIC-BLA or pIC-BLA projection neurons. **(K)** Fluorescence image of aIC-BLA projection neurons expressing mCherry (control virus). **(L)** Chemogenetic manipulations of aIC-BLA neuron do not alter NaCl-induced water intake (n=17 mCh, n=13 Gq, n=18 Gi, one-way ANOVA F(2,45)=0.8775, p=0.4228) while activation of pIC-BLA neurons strongly reduces NaCl-induced water intake (n=9 mCh, n=11 Gq, n=8 Gi, one-way ANOVA, F(2,25)=24.36, ****p<0.0001; Gq vs Gi Tukey’s post-hoc test ****p<0.0001). Data in **(D-G)** and **(L)** are shown as the mean ± SEM. For more relevant information, see **Figures S3, S4,** and **Table S1**.

Optic fibers were then implanted in the aIC or pIC (**Figures 3A, S3A, and S3B**) to record calcium signal in aIC-BLA or pIC-BLA projection neurons (**Figures 3B, S3A, and S3B**), respectively. Interestingly, aIC-BLA neurons displayed a decrease in neural activity at the onset of a licking event that lasted for several seconds (**Figures 3C-3E**), whereas pIC-BLA neurons showed an increase in activity at the onset of licking that persisted for the entire duration of licking bouts (**Figures 3C, 3F, and 3G**). Activity during late licks was similar to postlicking in mice recorded in both aIC-BLA and pIC-BLA projection neurons (**Figures S3C-S3F**). These results indicate that aIC-BLA and pIC-BLA neurons differentially respond to water intake, suggesting the two populations of neurons might have antagonistic control over drinking behavior.

### Posterior insula neurons targeting the BLA (pIC-BLA) negatively regulate water intake

To investigate the causal relationship between the neural activity of IC-BLA projection neurons and drinking behavior, we used a chemogenetic approach, expressing excitatory or inhibitory designer receptors exclusively activated by designer drugs (DREADDs) to manipulate neural activity [36–38]. The impact of chemogenetic manipulations was observed on water intake triggered by a systemic hypertonic NaCl administration as well as on basal levels of water intake (**Figures 3H and 3I**). To express DREADDs (Gq or Gi) in aIC-BLA or pIC-BLA projection neurons, we injected a retrograde AAV expressing the Cre recombinase into BLA, in combination with the injection of an AAV virus carrying Cre-dependent expression of the excitatory or inhibitory DREADDs in the aIC or pIC [36, 38] (**Figure 3J-K, S4A-B)**. Interestingly, activation of pIC-BLA, but not aIC-BLA, projection neurons strongly decreased water intake induced by hypertonic NaCl treatment and basal drinking, in comparison to the control group (**Figures 3L and S4C-G**). However, inhibition of aIC-BLA or pIC-BLA projection neurons did not alter water consumption (**Figures 3L and S4C-G**). Taken together, these experiments reveal that pIC-BLA neurons play an inhibitory role on water intake.

### Selective depolarization-induced suppression of excitation (DSE) at pIC-BLA terminals

The data obtained so far show that lack of CB_1_ expression in pIC neurons, as well as hyperactivation of pIC-BLA projections containing CB_1_ receptors, both reduce water intake. This suggests that CB_1_ receptors might negatively control pIC-BLA synaptic transmission. Stimulation of presynaptic CB_1_ receptors by endocannabinoids (eCB) retrogradely released from a depolarized postsynaptic neuron, suppresses neural transmission [26–29]. Thus, we hypothesized that CB_1_ receptors on axonal terminals of IC neurons in the BLA could express endocannabinoid-dependent depolarization-induced suppression of excitation (DSE). To test this idea, we used *ex vivo* optogenetically assisted circuit mapping combined with whole-cell patch clamp recordings in the BLA [39, 40]. We injected AAV carrying the gene coding for Channelrhodopsin-2 fused to the fluorescent protein mCherry under the control of the CaMKII promoter (AAV-CaMKII-ChR2-mCh) into the aIC or pIC of wild type mice (**Figures 4A-4C**). Four weeks later, we recorded optically-evoked monosynaptic excitatory postsynaptic currents (oEPSCs) in BLA neurons induced by blue light photostimulation of IC-BLA axon terminals. Remarkably, 5 seconds depolarization of the postsynaptic BLA neurons induced DSE of pIC-BLA photocurrents in 8 out of 15 neurons recorded in wild type animals (**Figures 4D, 4F, and S4H**). Interestingly, only 1 out of 10 cells expressed DSE in global *CB_1_*-KO mice. The proportion of cells expressing DSE is more than five times higher in wild type mice compared to *CB_1_*-KO mice (**Figures 4F and S4I**), suggesting that CB_1_ receptors in pIC-BLA axon terminals contribute to the regulation of these synapses through DSE. Interestingly, no DSE was observed when oEPSCs were obtained from aIC-BLA axon terminals (**Figure 4G**). In addition, stimulation of cannabinoid receptors by a synthetic agonist, WIN 55,212-2 (WIN), produced a reduction of oEPSCs in wild type mice but not *CB_1_*-KO controls (**Figure S4J**). Therefore, these data reveal that CB_1_ receptors on pIC-BLA, but not aIC-BLA, axon terminals are stimulated by endocannabinoids to inhibit transient excitatory neurotransmission.

**Figure 4.**
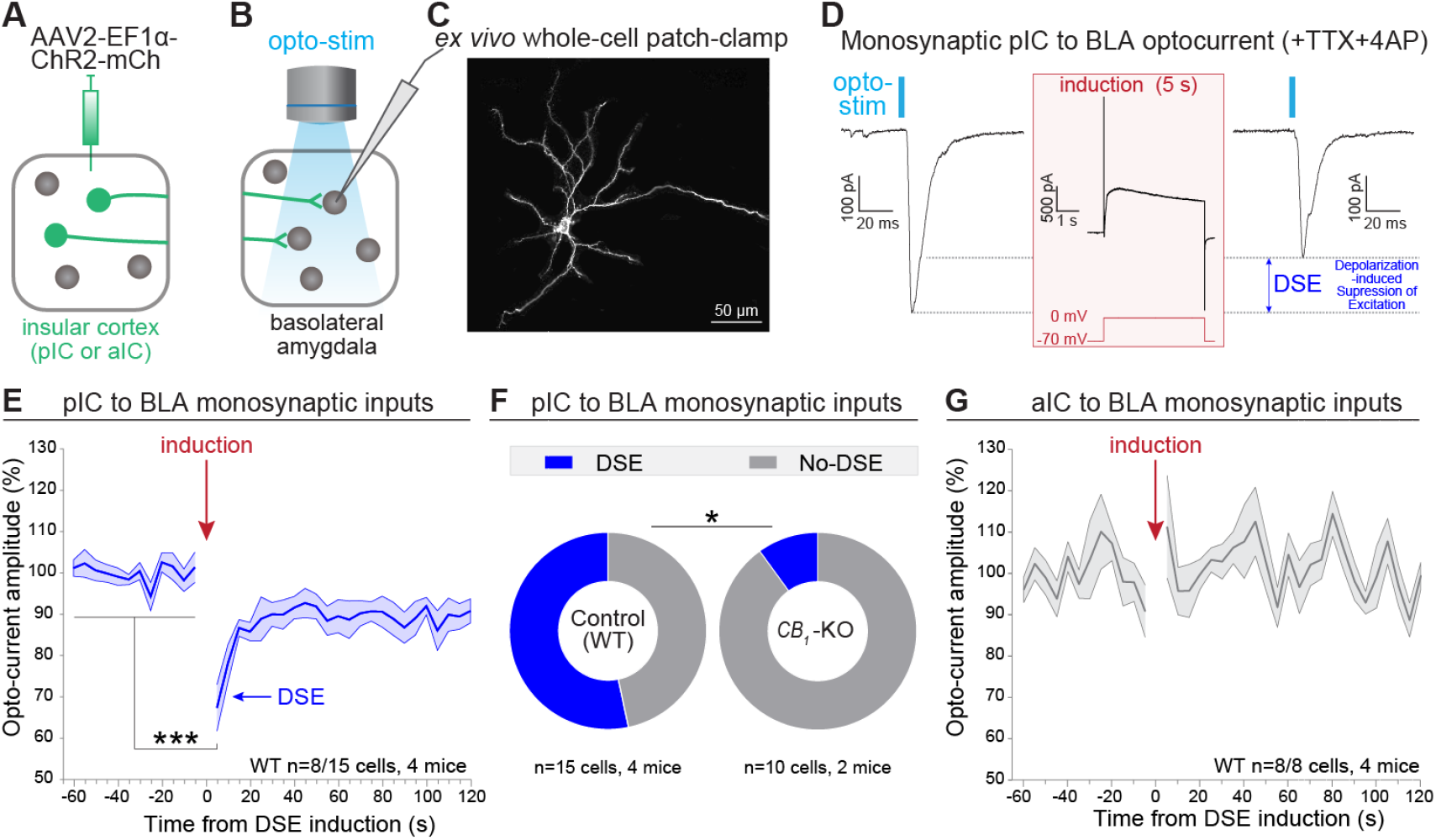
Selective depolarization-induced suppression of excitation (DSE) at pIC-BLA terminals. **(A)** ChR2 was selectively expressed in pIC or aIC neurons. **(B)** Diagram of *ex vivo* whole-cell patch-clamp recording of BLA principal neurons coupled with optogenetic stimulation of insular inputs. **(C)** Confocal image of a representative neuron recorded in the BLA. **(D)** Representative monosynaptic currents recorded in a BLA neuron during optogenetic stimulation of pIC axons (473 nm, 5 ms), before and after DSE induction (induction protocol: 0 mV, 5 s). Monosynaptic currents were isolated thanks to the presence of 1 μM TTX and 100 μM 4AP. **(E)** Time course of pIC to BLA current amplitude, average over neurons that exhibit DSE (two-tailed paired *t*-test, n=8/15 cells from 4 mice; ***p=0.0007). **(F)** The proportion of cells expressing DSE is higher in WT compared to *CB_1_*-KO mice (chi-square, *p=0.04). **(G)** Time course of aIC to BLA monosynaptic currents (two-tailed paired *t*-test, n=8/8 from 4 mice; p=0.396). Data in **(E)** and **(G)** are shown as the mean ± SEM. For more relevant information, see **Figure S3**.

## DISCUSSION

This study shows that both activation of pIC-BLA projection neurons, and removal of CB_1_ receptor in the pIC, negatively regulate water intake. As we found that pIC-BLA synapses express CB_1_-dependent synaptic plasticity, our findings suggest that CB_1_ receptors within the pIC-BLA pathway control water intake.

After observing that neural activity of excitatory neurons in both aIC and pIC increases in response to water consumption in freely moving mice, we found that removal of CB_1_ receptors, only in pIC cells, suppresses water intake. We then found that pIC outputs in the BLA, but not the CeA, express CB_1_ receptors, and that pIC-BLA synapses selectively express CB_1_-dependent depolarized suppression of excitation (DSE). Thus, we hypothesized that the behavioral impact of insular CB_1_ knock-out is due to the loss of inhibition of excitatory neurotransmission from pIC axonal terminals to BLA neurons. Consistently, we identified that the activity of pIC-BLA projection neurons is selectively increased in response to water licking and that chemogenetic stimulation of pIC-BLA projection neurons inhibits water intake. Altogether, our results support a model where pIC-BLA activation encodes water satiety and CB_1_ receptors located on pIC axonal terminals in the BLA positively control water intake through inhibition of pIC-BLA neurotransmission.

### Coding and control of water licking in projection neurons of the anterior and posterior insula

Thirst has been positively correlated with IC neural activity in both humans and rodents [7–9]. However, a recent study reported that part of the insula neurons increase while others decrease their firing rate when physiological state transfers from thirst to sated state, suggesting that subpopulations of IC neurons oppositely encode internal water balance conditions [11]. Here, we found that at a neural population level, in freely moving mice, the activity of excitatory neurons of both aIC and pIC increases in response to water drinking. Interestingly, a previous study showed that optogenetic stimulation of aIC and pIC projection neurons during water consumption respectively increases and decreases lick numbers [41]. Together with our results, these data show that aIC and pIC projection neurons are both activated in response to licking water but oppositely control water consumption, and suggest that pIC glutamatergic neurons encode a water satiety signal.

### Differential role of CB_1_ receptors in different cortical areas in the control of water intake

Previous studies have reported that global deletion or blockade of CB_1_ receptors impairs drinking behavior induced by hypertonic NaCl treatment [15, 16]. Indeed, mice without expression of CB_1_ receptors (Stop-*CB_1_* mice) drink less water than mice with global re-expression of CB_1_ receptors (*CB_1_*-RS mice) in response to hypertonic NaCl treatment [16]. In this study, deletion of CB_1_ receptors in cells of the entire IC, or the pIC, but not aIC, reduced water intake, suggesting that CB_1_ receptors in pIC cells are necessary to guarantee a physiological amount of water intake. However, selective re-expression of CB_1_ receptors in IC cells of Stop-*CB_1_* mice did not affect drinking behavior compared to control mice [16]. Thus, CB_1_ receptors in the IC are necessary to allow physiological amounts of water intake, but are not sufficient to rescue alteration of water intake induced by global deletion of CB_1_ receptor expression. Conversely, re-expression of CB_1_ in the anterior cingulate cortex (ACC) neurons in Stop-*CB_1_* mice rescued physiological water intake [16]. Altogether these results suggest that CB_1_ receptors in the pIC and ACC are both involved but play different roles in the control of water intake.

### Dichotomy of insula CB_1_ receptors expression on IC inputs to amygdala territories

Both aIC and pIC neurons send projections to the BLA [30–32]. A recent study reported that CB_1_ receptors are expressed in IC glutamatergic neurons [42], but the distribution of CB_1_ receptors in axonal terminals of IC neurons is unclear. Interestingly, our targeted histological analyses revealed that CB_1_ receptors of IC neurons are intensively located on BLA but not CeA axon terminals. In the BLA, CB_1_ receptors were shown to be located at both excitatory and inhibitory synapses, and activation of those receptors decreased glutamatergic and GABAergic synaptic transmission [43–45]. CB_1_ mRNAs colocalize with cholecystokinin (CCK) mRNAs in the BLA [25], and CB_1_ receptors/CCK form symmetrical synapses in the BLA, suggesting that CB_1_ receptors are present in CCK interneurons and locally form inhibitory synapses [43, 46]. However, a recent study also reported that most CCK neurons in the BLA are glutamatergic neurons, with only a small fraction of CCK-positive cells being GABAeric neurons [47]. Despite the knowledge of CB_1_ receptors in the BLA, little was known on the potential source of CB_1_ receptors from long-range glutamatergic inputs. This study is the first to show that CB_1_ receptors are located on pIC-BLA synapses, while our previous study showed they are also present on ACC-BLA terminals [16].

### Antagonistic activity of aIC-BLA and pIC-BLA projection neurons during water intake

We revealed that the global activity of aIC-BLA projection neurons decreases in response to water licking, whereas the activity of pIC-BLA projection neurons increases. This suggests that these two neuronal populations oppositely code drinking behavior. A recent study reported that aIC-BLA projection neurons decrease activity when mice lick sucrose, whereas aversive stimuli including mild foot shock and tail suspension produce an increase in activity of these neurons [30]. This suggests that aIC-BLA projection neurons code for negative valence, which in the context of water intake could be associated with thirst. As pIC-BLA neurons have an opposite role, we suggest that they might likely participate in the encoding of water satiety.

### State and environment dependent roles of aIC-BLA projection neurons in water drinking

Chemogenetic manipulation of aIC-BLA projection neuron did not alter water intake, which was consistent with previous findings [14], but also unexpected in light of a previous study showing that optogenetic stimulation of aIC-BLA axon terminals could increase water licking [13]. These apparently discrepant results are possibly due to differences in the experimental approaches. For instance, whereas Wang et al. measured conditioned responses in head-restrained mice, where water was announced by a light cue, and available for only 5 seconds following the cue, in the present study water intake was triggered by a hypertonic injection of NaCl, and mice were freely moving in their home cage [13]. Moreover, the stimulation approaches are also different, as Wang et al., employed optogenetics, with a behavioral closed-loop stimulation protocol initiated by licking. Instead, we used Gq-DREADD expression to produce longer activation over the entire drinking test. Thus, the difference in environmental and sensory factors, as well as duration of manipulation might produce brain states driving different behaviors.

### Negative control of pIC-BLA projection neurons on water drinking

Using chemogenetic manipulation, we identified that activation of pIC-BLA, but not aIC-BLA, projection neurons reduces drinking induced by hypertonic NaCl treatment. This is consistent with the hypothesis that activity in this neural population encodes water satiety. However, inhibition of these neurons did not promote water intake, potentially due to a ceiling effect. To test this possibility, we investigated how pIC-BLA neurons regulate basal drinking. In this experiment, mice did not receive any treatments and had *ad libitum* water access. In these conditions, activation of pIC-BLA projection neurons also inhibited drinking, however, inhibition of these neurons still did not alter water drinking. A recent study showed that stimulation of pIC excitatory neurons promotes context-induced feeding [48], suggesting that the stimulation of these neurons does not induce malaise and suppresses general intake behavior. Altogether, these data indicate that pIC-BLA projection neurons exert a negative, but not positive, control on water intake, compatible with the encoding of a water satiety signal.

### Selective CB_1_-dependent DSE in pIC-BLA pathway

Previous studies showed that depolarization-initiated cannabinoid release suppresses synaptic transmission through stimulation of CB_1_ receptors in the BLA [33, 34, 49]. However, the cell types expressing the CB_1_ receptors and mediating DSE remained unknown. In this study we identified for the first time that CB_1_ receptors specifically in the pIC-BLA pathway mediate DSE. Indeed, we found that CB_1_-dependent DSE is present in the pIC-BLA pathway, but almost absent in the aIC-BLA pathway, which confirms that CB_1_ receptors at glutamatergic pIC-BLA synapses are functional.

### Conclusions

This study dissects the role of subpopulations of insula neurons in the control and coding of water intake. The data reveal that the activity of pIC and pIC-BLA projection neurons increases in response to water intake, and that chemogenetic stimulation of pIC-BLA neurons negatively regulates drinking behavior, and potentially encodes a water satiety signal. At the molecular level, CB_1_ receptors in the pIC also control water intake, and can regulate synaptic transmission from the pIC to the BLA, through retrograde endocannabinoids-induced inhibition of excitatory neurotransmission. Altogether, this work provides a mechanistic model of insular cortex control of water intake to maintain physiological homeostasis.

## Supporting information

Zhao_et_al_Supplementary_video1

Zhao_et_al_Supplementary_video1

## SUPPLEMENTARY FIGURE TITLES AND LEGENDS

**Figure S1.**
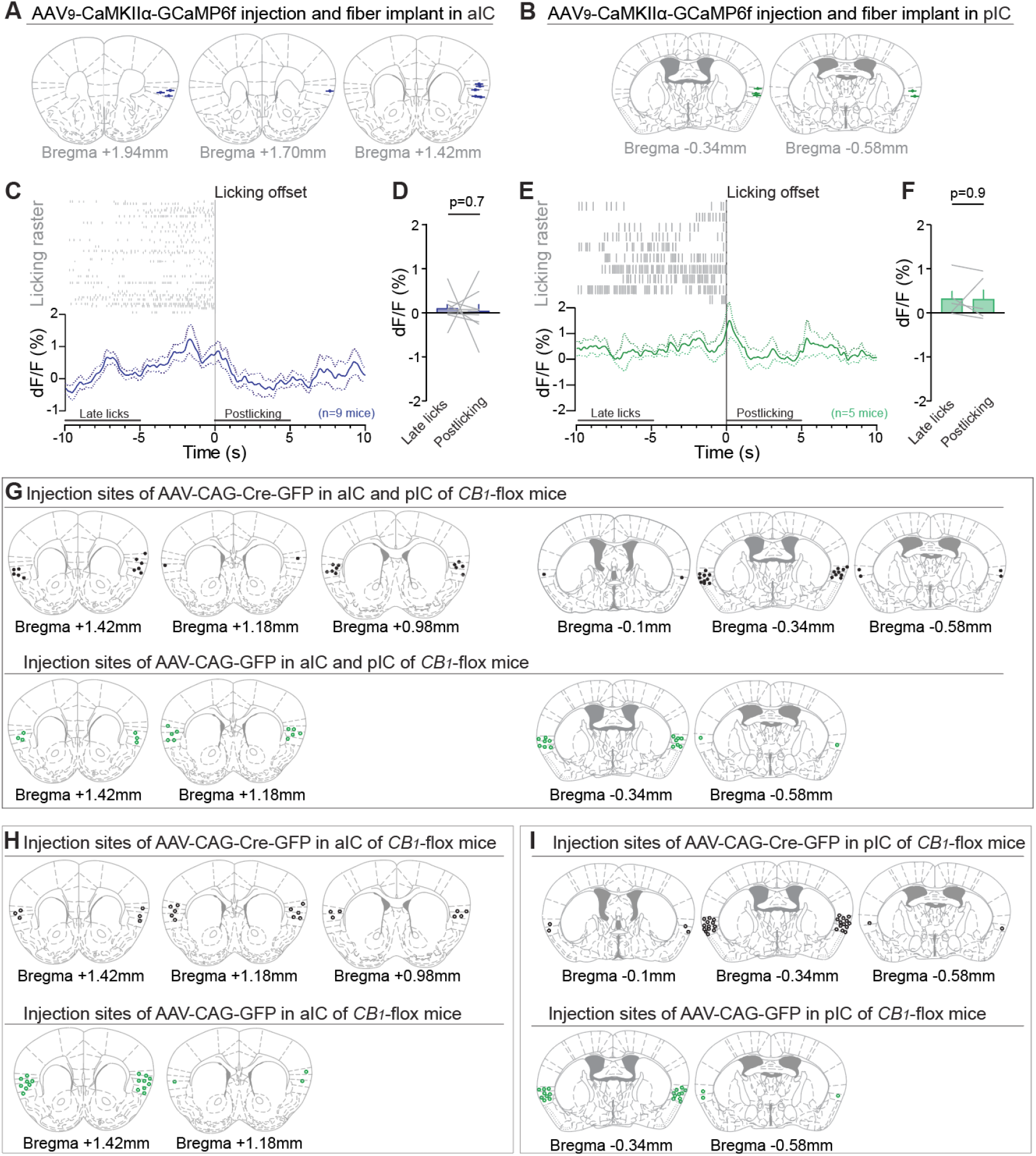
Histological verification of GCaMP6f expression and location of fiber implantation in aIC or pIC, and Cre recombinase or GFP expression in IC of *CB_1_*-flox mice. **(A** and **B)** Location of viral injection sites and fiber implants in the aIC **(A)** or pIC **(B)**. **(C)** Top, licking raster of drinking bouts. Bottom, an average of aIC calcium signal of drinking bouts. Signal of each bout is aligned to last lick, the traces from −10 to −5 s and from 0 to 5 s are considered as late licks and postlicking signals respectively. dF/F represents the fluorescent changes from the mean level of the entire recording time series. Dashed lines are mean ± SEM. **(D)** Mean of calcium signal in aIC during Late licks and Postlicking phases. Neural activity during Late licks phase in aIC is similar with the activity during Postlicking phase (two-tailed paired *t*-test, n=9, p=0.7,). **(E** and **F)** Same as **(C)** and **(D)**. Neural activity during Late licks phase in pIC is similar with the activity during Postlicking phase (Two-tailed paired *t*-test, n=5, p=0.9). **(G)** Top, location of viral injection sites of Cre-expressing virus (AAV-CAG-Cre-GFP) in the aIC and pIC of CB_1_-flox mice (IC-CB_1_-KO mice); one mouse was excluded because of no Cre expression. Bottom, location of viral injection sites of GFP-expressing virus (AAV-CAG-GFP) in aIC and pIC of *CB_1_*-flox mice (control mice). **(H)** Top, location of viral injection sites of Cre recombinase in aIC of *CB_1_*-KO mice; 5 mice were excluded because of wrong targets or no Cre expression. Bottom, location of viral injection sites of GFP in aIC of control mice. **(I)** Top, location of viral injection sites of Cre recombinase in pIC of *CB_1_*-KO mice; one mouse was excluded because of no Cre expression. Bottom, location of viral injection sites of GFP in pIC of control mice.

**Figure S2.**
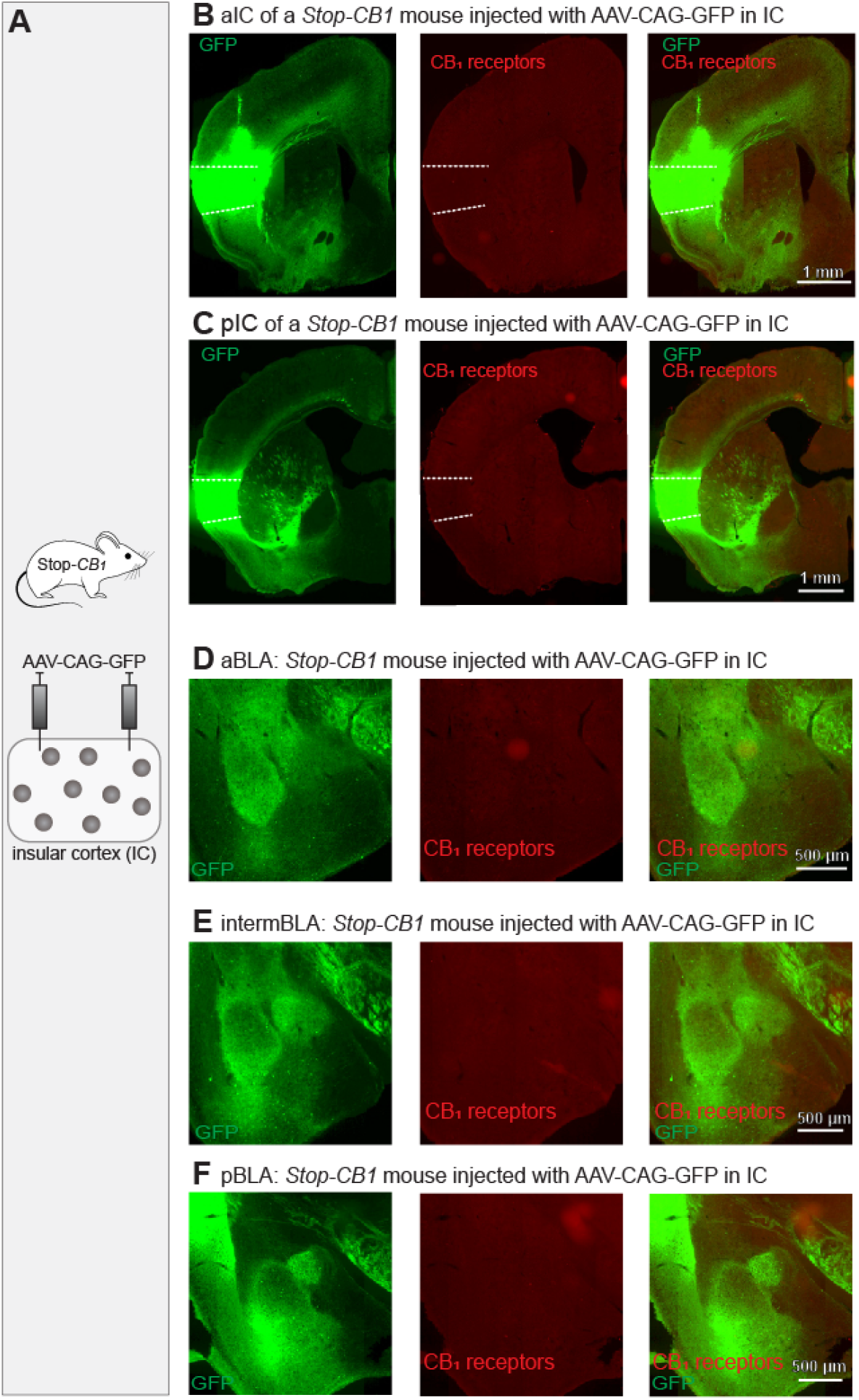
CB_1_ receptors are absent in the IC and amygdala of Stop-*CB_1_* control mice. **(A)** Schematic presentation of viral expression of GFP in IC of Stop-*CB_1_* mice. **(B-F)** CB_1_ receptors are absent in aIC **(B)**, pIC **(C)**, aBLA **(D)**, intermBLA **(E)**, pBLA **(F)** in Stop-*CB_1_* control mice. Scale bar, 1 mm.

**Figure S3.**
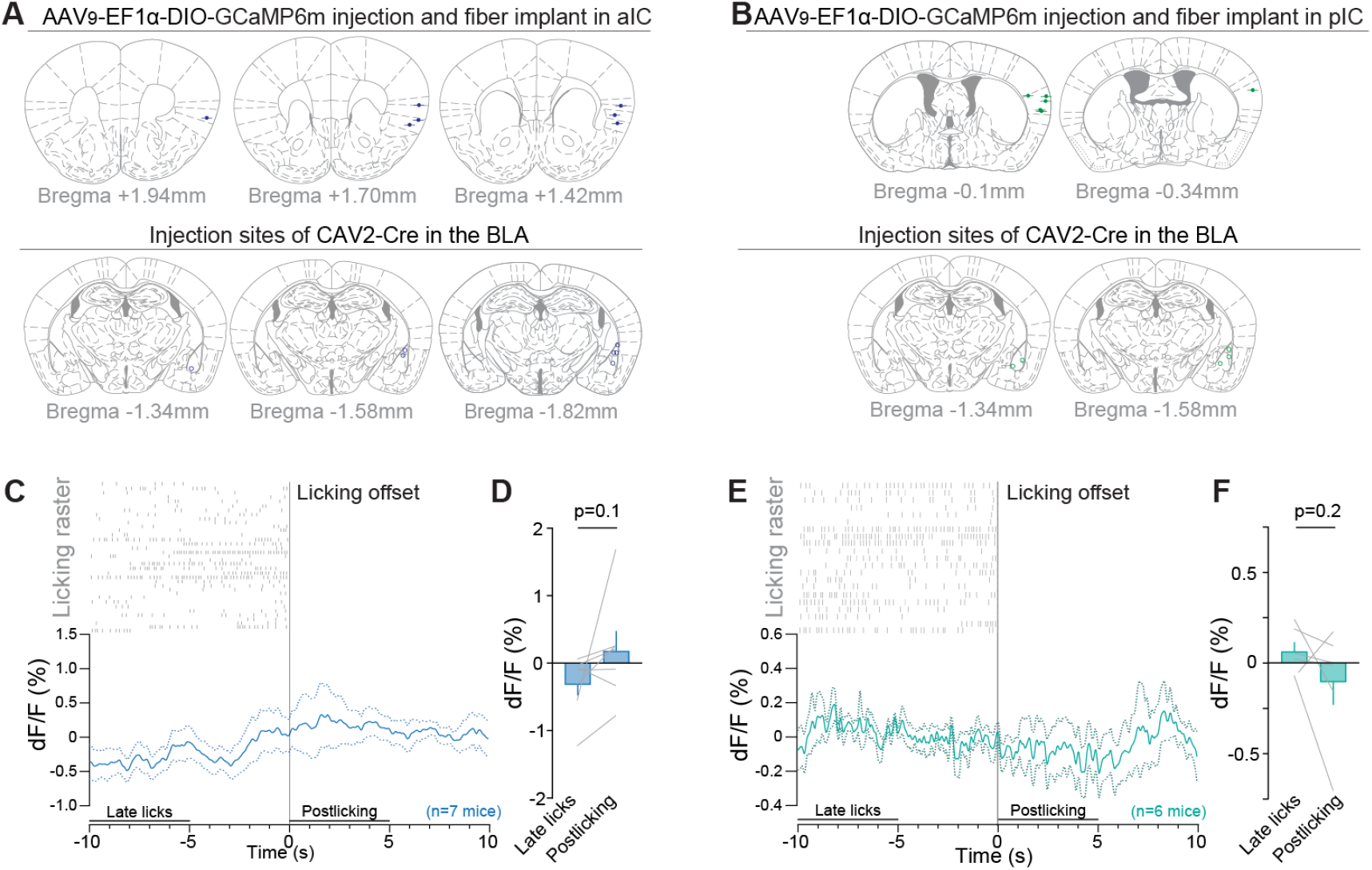
Histology and licking offset responses for IC-BLA fiber photometry recordings. (aIC-BLA or pIC-BLA). **(A)** Location of the anterograde viral vector injection, and fiber implant in the aIC (top), and of the retrograde virus injection in the BLA (bottom) in mice expressing GCaMP6m in aIC-BLA projection neurons. **(B)** Location of the injection of the anterograde viral vector and fiber implant in the pIC (top), and of the retrograde vector injection in the BLA (bottom) in mice expressing GCaMP6m in pIC-BLA neurons. **(C)** Lick raster of drinking bouts above the average calcium signal of aIC-BLA neurons recorded during 36 drinking bouts. The signal of each bout is aligned to the last lick of each bout. The activity from −10 to −5 s and from 0 to 5 s is considered as Late licks and Postlicking signals respectively. dF/F represents the fluorescent changes from the mean level of the entire recording time series. Dashed lines represent mean ± SEM. **(D)** Neural activity of aIC-BLA projection neurons tends to be higher in the ‘Postlicking phase’ compared to the ‘Late licks phase’ (two-tailed paired *t*-test, n=7, p=0.1094). **(E)** Lick raster of drinking bouts above the average calcium signal of pIC-BLA neurons recorded during 21 drinking bouts. The signal of each bout is aligned to the last lick of each bout. Dashed lines represent mean ± SEM. **(F)** In pIC-BLA neurons, the calcium signal is similar in the Late licks and Postlicking phases (two-tailed paired *t*-test, n=6, p=0.2390).

**Figure S4.**
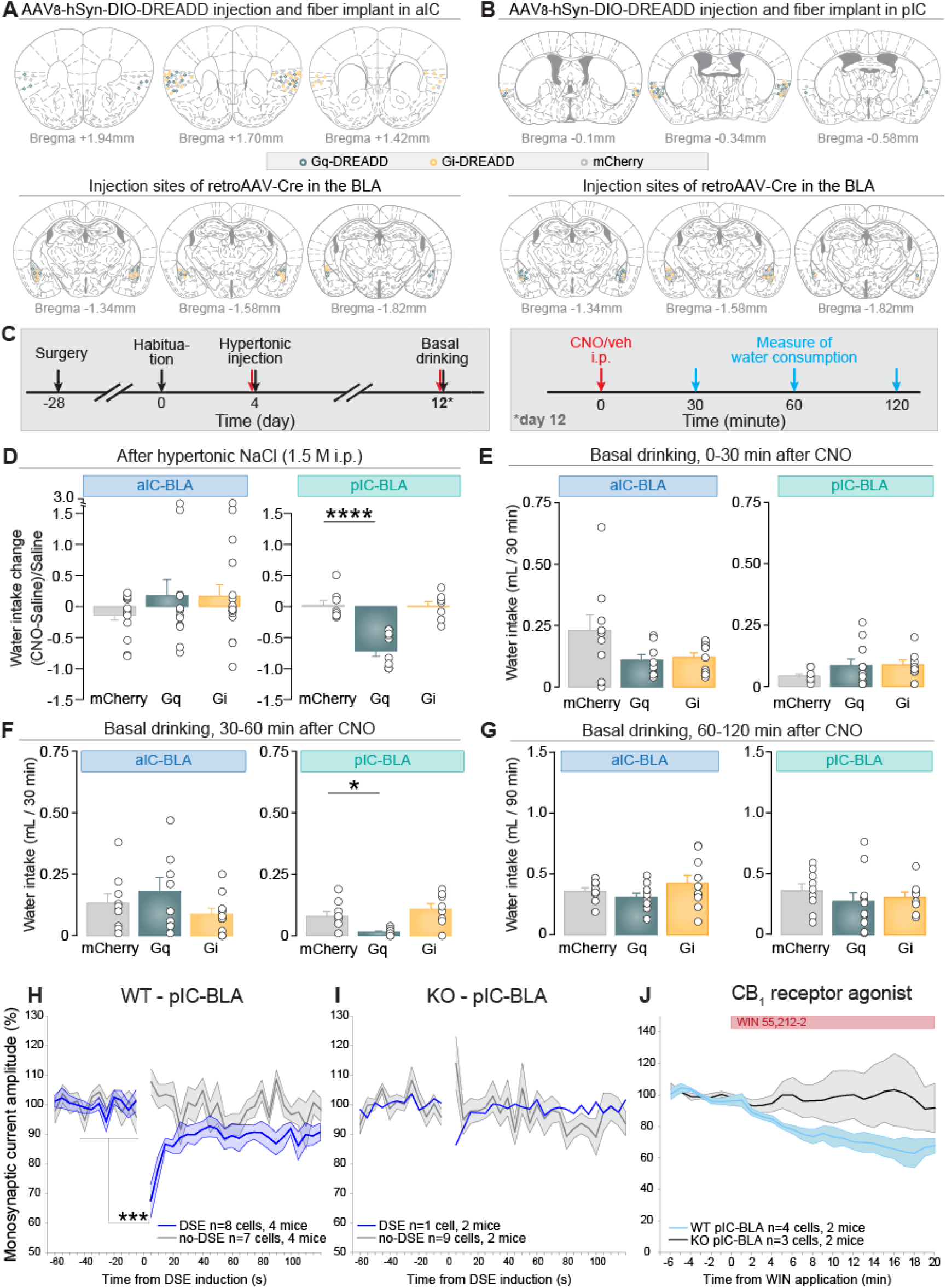
Histology and drinking behavior in DREADDs or mCherry expression in IC-BLA neurons (aIC-BLA or pIC-BLA), and averaged monosynaptic pIC-BLA current time course recorded in BLA neurons. <u>**(A-B)** Hostological verification</u> **(A)** Location of viral injection sites in the aIC (top) and BLA (bottom) of mice expressing the Gq-DREADD (aquamarine circle), Gi -DREADD (yellow circle) and mCherry (grey circle) in aIC-BLA projection neurons. **(B)** Location of viral injection sites in the pIC (top) and BLA (bottom) for mice expressing the Gq-DREADD (aquamarine circle), Gi-DREADD (yellow circle) and mCherry (grey circle) in pIC-BLA neurons. 5 aIC-BLA-Gq-DREADD mice were excluded because of wrong targets or no DREADD-Gq expression. <u>**(C-G)** Changes in drinking behaviors induced by CNO injections</u> **(C)** Experimental design of the chemogenetic experiment and basal water intake measurements. **(D)** Change of drinking behavior induced by CNO injection relative to saline injection in each mouse. pIC-BLA-Gq mice drink less after CNO injection than after saline injection, compared to pIC-BLA-control mice (n=11 pIC-BLA-Gq mice, n=9 pIC-BLA-control mice, one-way ANOVA, ****p<0.0001). **(E-G)** Basal water intake during 0-30, 30-60, and 60-120 minutes after CNO injection. **(E)** Basal water intake in the first 30 minutes of the test (0-30 minutes) after CNO injection. **(F)** pIC-BLA-Gq mice drink less water between 30 and 60 minutes compared to pIC-BLA-control mice (n=10 pIC-BLA-Gq mice, n=8 pIC-BL-Gi mice, n=9 pIC-BLA control mice, one-way ANOVA test, F(2,24)=8.859, **p=0.0013, Gq vs control Tukey’s post-hoc test *p=0.02). **(G)** Basal water intake between 60 to 120 minutes of the test (60-120 minutes) after CNO injection. For more relevant information, see **Table S1**. <u>**(H-J)** pIC to BLA inputs responsiveness to endogenous and exogenous cannabinoids</u>. **(H)** Time course of pIC to BLA inputs recorded from neurons that underwent DSE (two-tailed paired *t*-test, n=8/15 cells from 4 mice, ***p=0.0007) and no DSE (two-tailed paired *t*-test, n=7/15 cells from 4 mice, p=0.125) in WT mice. **(I)** Time course of pIC to BLA inputs recorded from neurons that displayed DSE (n=1/10 cells from 2 mice) and no DSE (two-tailed paired *t*-test, n=9/10 cells from 2 mice, p=0.161) in *CB_1_*-KO mice. **(J)** pIC to BLA inputs responsiveness to 5 μM WIN 55,212-2 recorded from WT (two-tailed paired *t*-test, n=4 cells from 2 mice, p=0.016) and *CB_1_*-KO mice (two-tailed paired *t*-test, n=3 cells from 2 mice, p=0.885).

## STAR ⋆ METHODS

### RESOURCE AVAILABILITY

#### Lead Contact

Further information and requests should be directed to and will be fulfilled by the Lead Contact (Anna Beyeler).

#### Materials Availability

This study did not generate new unique reagents. Materials used here are available from the Lead Contact upon reasonable request.

#### Data and Code Availability

Raw data and code supporting the current study (Figures 1-4 and S1-S4) have been deposited to Mendeley Data: doi: 10.17632/v3b2fbrd24.1

### EXPERIMENTAL MODEL AND SUBJECT DETAILS

All experiments were approved by the ethical Committee on Animal Health and Care of INSERM and the University of Bordeaux, and by the French Ministry of Agriculture and Forestry (authorization numbers 15493 and 12411). Maximal efforts were made to reduce the suffering and the number of mice used. Drinking behavior triggered by systemic hypertonic NaCl treatment was performed during the light phase, fiber photometry recordings and basal drinking measurements were done in dark phase. Animals were kept in individual cages under standard conditions in a day/night cycle of 12/12 hours (lights on at 7 am) for drinking behavioral test. For the fiber photometry recordings, mice were grouped-housed (3-6 mice) and kept in a reversed light-dark cycle. Male wild-type C57BL/6 mice were purchased from Janvier (France). All mutant mice were generated and identified in previous studies, e.g. *CB_1_*-Flox mice; the Stop-*CB_1_* mice (lack of CB_1_); global CB_1_ knockout (*CB_1_*-KO) mice [22, 33, 50]. All the mice used in this study were 7-10 weeks old at the beginning of the experiments and all the data was obtained by experimenters blind to viral expression or genetic conditions.

### METHOD DETAILS

#### Surgery and viral administration

Mice were anesthetized by isoflurane (5% for the induction and 2% during the surgery) and placed on a stereotaxic apparatus (Model 900, KOPF instruments, CA, USA) with a mouse adaptor and lateral ear bars. For viral vectors delivery, AAV vectors were loaded in a glass pipette and infused by a pump (UMP3-1, World Precision Instruments, FL, USA). The injection coordinates in anteroposterior (AP) / mediolateral (ML) / dorsoventral (DV) from Bregma, were in mm: for the aIC +1.7/±3.0/-3.5, for the pIC −0.3/±3.7~4.0/-4.0 (200~300 nL per injection site, 100 nL/min), for the BLA −1.6/±3.3/-4.9 (150 nL, 100 nL/min). The coordinates used were decided according to the mouse brain atlas [51].

<u>For fiber photometry experiments</u> targeting all excitatory neurons, 300 nL of AAV9-CaMKIIα-GCaMP6f (Addgene 100834, >1×10^13^ vg/mL) was injected into the right aIC or right pIC, and an optical fiber (400 μm diameter, 0.39 numerical aperture (NA)) was implanted 50 μm above the viral injection site. To express GCaMP6m in aIC-BLA or pIC-BLA projection neurons, 300 nL of CAV2-Cre vector (IGMM, 1.12×10^13^ pp/mL) was injected into the right BLA, in combination with an injection of 300 nL of AAV9/2-EF1α-DIO-GCaMP6m vector (Stanford University, GVVC-AAV-92, >1×10^12^ GC/mL) into the right aIC or pIC. Directly after viral injections, an optical fiber (400 μm diameter, 0.39 NA) was implanted 50 μm above the viral injection site in the aIC or pIC.

<u>For deletion of CB_1_ receptors</u> in the IC, 200 nL of AAV1/2-CAG-GFP (custom made, 5.18×10^10^ vg/mL) or AAV1/2-CAG-Cre-GFP (custom made, 4.2×10^10^ vg/mL; for control mice) were injected into the aIC, the pIC, or both (entire IC) at 100 nL/min in *CB_1_*-Flox mice.

<u>For re-expression of CB_1_ receptors</u> in IC, 200 nL of AAV1/2-CAG-Cre-GFP (custom made, 4.2×10^10^ vg/ml) was injected into the entire insula (IC) including both aIC and pIC at 100 nL/min in Stop-*CB_1_* mice.

<u>To map CB_1_ mRNA in IC-BLA neurons</u>, 150 nL of retroAAV2-hSyn1-chI-EBFP produced by the Zurich Neuroscience Center (ZNZ) Viral Vector Facility (VVF) was injected in the BLA (ZNZ-VVF v140-retro, 4.1×10^12^ vg/ml).

<u>For manipulating the activity of IC-BLA neurons with DREADDs</u>, 200 nL of AAV8-hSyn-DIO-hM3D(Gq)-mCherry (Addgene 443618, 2.2×10^13^ GC/mL), AAV8-hSyn-DIO-hM4D(Gi)-mCherry (Addgene 50475, 2.1×10^13^ GC/mL), or AAV8-hSyn-DIO-mCherry (Addgene 50459, 2.3×10^13^ GC/mL) was injected bilateraly into the aIC or pIC, in combination with the injection of 150 nL of retroAAV2-hSyn1-chI-iCre-EBFP (Addgene #25493, ZNZ VVF v148, 6.7×10^12^ vg/mL) into the BLA.

<u>For *ex vivo* whole-cell patch-clamp</u> recording experiments, 200 nL of AAV2/2-hSyn1-hChR2(H134R)-mCherry (Addgene #58880, ZNZ-VVF v124, 3.3×10^12^ vg/ml) was injected into the aIC or pIC of WT or *CB_1_*-KO mice.

#### *In vivo* calcium imaging by fiber photometry system

Fiber photometry was performed as in previous studies [30, 52, 53]. A 470 nm LED was used to excite GCaMP6 in a calcium-dependent manner and a 405 nm LED was used as an isosbestic wavelength to excite GCaMP in a calcium-independent manner (LEDs from Thorlabs). Acquired photometry data were processed with custom-written codes in MATLAB. To calculate ΔF/F time series, a linear fit was applied to the 405 nm signals and aligned to the 470 nm signals. The fitted 405 nm signal was subtracted from the 470 channel, and then divided by the fitted 470 nm signal to obtain ΔF/F values. Photometry signals were then extracted from 10 seconds before to 10 seconds after each licking bout of continuous licks of 5 seconds or more and averaged across licking bouts of different animals.

#### Water intake assays

For the experiments with CB_1_ receptors deletion in the insula, the amount of water intake was measured 30, 60, and 120 minutes after the intraperitoneal (i.p.) injection of 1.0 M sodium chloride (NaCl, VWRV0241, 10 mL/kg of body weight). For the DREADD experiments, the amount of water intake was measured 30, 60, and 120 minutes after the i.p. injection of 1.5 M sodium chloride (NaCl, VWRV0241, 10 mL/kg of body weight), and Clozapine-N-oxide (CNO, 2 mg/kg, Tocris Bioscience) or the control saline were injected 30 minutes before the water intake test (**Figure 3I**). To control that mice were drinking normally before the manipulations, the daily water intake and body weight of each mouse was monitored daily during the whole period of experiments.

#### Immunohistochemistry (IHC)

After the behavioral experiments, mice were anesthetized with pentobarbital (Exagon, 400 mg/kg body weight), transcardially perfused first with the phosphate-buffered solution (PBS, 0.1M, pH 7.4) and then fixed by 4% formaldehyde (Sigma-Aldrich, HT501128, [21, 23]. Serial brain coronal sections were cut at 40 μm and collected in PBS at room temperature (RT). Sections were permeabilized in a blocking solution of 4% donkey serum, 0.3% Triton X-100 and 0.02% sodium azide prepared in PBS for 1 hour at RT. For the CB_1_ immunohistochemistry, free-floating sections were incubated with goat CB_1_ receptors polyclonal primary antibodies (CB_1_-Go-Af450-1; 1:2000, Frontier Science Co. ShinKO-nishi, Ishikari, Hokkaido, Japan) for 48 hours at 4°C. The antibody was prepared in the blocking solution. After three washes, the sections were incubated with a secondary antibody anti-goat Alexa Fluor 555 (A21432, 1:500, Fisher Scientific) for 2 hours at RT and then washed in PBS at RT. Images of these sections were taken by a Nanozoomer microscope (Hamamatsu, Japan) and Leica SP8 confocal microscope (Leica, Germany) and processed using Fiji (Image J,NIH).

#### Fluorescent *in situ* hybridization (FISH) coupled with immunohistochemistry

The detailed procedure referred to previous publications [23–25]. Mice were sacrificed by cervical dislocation, then their brains were rapidly extracted and placed on dry ice. The frozen brains were stored at 80°C for sections by a cryostat (40 μm, CM1950, Leica). Coordinates of selected pIC sections were taken from Bregman +0.2 to −0.8 mm. For the probes, fluorescein (FITC)-labeled riboprobes against mouse CB_1_ receptor was made by our lab [25]. After hybridization overnight at 60°C with the mixture of probes, the slides were washed with different stringency wash buffers at 65°C. Then, the slides were blocked with a blocking buffer prepared according to the manufacturer’s protocol. Anti-FITC antibodies conjugated to horseradish peroxidase (HRP) (Roche; 1:2000) were applied 2 hours at RT or overnight at 4°C to detect respectively CB_1_-FITC probes. Probes hybridization was revealed by a tyramide signal amplification (TSA) reaction using Cyanine 3-labeled tyramide (Perkin Elmer; 1:100 for 10 minutes) to detect FITC-conjugated tyramide (Perkin Elmer; 1:80 for 12 minutes) to amplify the signal of CB_1_ as red color. For the amplification of blue fluorescent protein signals, immunohistochemistry (IHC) was applied by using rabbit anti-green fluorescent protein (GFP) antibody (A11122, 1:1000, Fisher Scientific) here. After incubation overnight in the primary antibody at 4°C, the sections were incubated with a secondary antibody anti-rabbit Alexa Fluor 488 (A11034, 1:500, Fisher Scientific) for 2 hours at RT and then washed in PBS at RT. After FISH and IHC, the slides were incubated in 4’,6-diamidino-2-phenylindole (DAPI; 1:20,000; Fisher Scientific) for 5 minutes. Then, slides were mounted with coverslips, visualized by Leica SP8 confocal microscope (Leica, Germany), and images were processed with Image J (NIH, see quantification and statistical analysis section).

#### Depolarization-induced suppression of excitation (DSE)

##### Brain tissue preparation

About 12-14 weeks after viral injection in the insular cortex, mice were anesthetized with 40 mg/kg pentobarbital and perfused transcardially, after clamping the abdominal aorta, with 20 mL of modified artificial cerebrospinal fluid (ACSF, at ~4°C) containing (in mM): 75 sucrose, 87 NaCl, 2.5 KCl, 1.3 NaH2PO4, 7 MgCl2, 0.5 CaCl2, 25 NaHCO3 and 5 ascorbic acid. The brain was then extracted and glued on the platform of a semiautomatic vibrating blade microtome (VT1200; Leica). The platform was then placed in the slicing chamber containing modified ACSF at 4°C. Coronal sections of 300 μm containing the insula or containing BLA/CeA were collected in a holding chamber filled with ACSF saturated with 95% O2 and 5% CO2, containing (in mM): 126 NaCl, 2.5 KCl, 1.25 NaH2PO4, 1.0 MgCl2, 2.4 CaCl2, 26 NaHCO3, 10 glucose. Recordings were started one hour after slicing and the temperature was maintained between 31–33°C both in the holding chamber and during the recordings. All viral injection sites in the IC were checked and imaged with the microscope (BX51, Olympus).

##### Whole-cell patch-clamp recording

Recordings were made from visually identified neurons in the BLA (BX51, Olympus, infrared illumination). Whole-cell recordings were acquired using glass microelectrodes (7-9 MΩ) filled with a solution containing (in mM): 120 cesium methansulphonate, 20 HEPES, 0.4 EGTA, 2.8 NaCl, 5 tetraethylammonium chloride, 2.5 MgATP, 0.25 NaGTP, 8 biocytin and 2 Alexa Fluor-350 (pH 7.25-7.4, 280-290 milliosmol). All recordings were made using a Multiclamp 700B amplifier (Molecular Devices). Analog signals were low-pass filtered at 1 kHz and digitized at 10 kHz using a Digidata 1550B and pClamp10 software (Molecular Devices). ACSF and drugs were applied to the slice via a peristaltic pump (Minipuls3, Gilson) at 2 mL/min.

ChR2 was activated using a LED light source (470 nm, CoolLED p4000) and the average power to trigger a light response in the patched neurons was 0.17±0.06 mW/mm^2^. To test if the response of ChR2 terminals activation was monosynaptic sodium channel blocker tetrodotoxin (TTX, 1 μM) was perfused in combination with the potassium channel blocker 4-aminopyridine (4AP, 100 μM) applied to facilitate glutamate release from synaptic terminals [39, 40].

The location of all recorded neurons was checked after the recording and only the cells located in the BLA were kept for further analysis. Amongst the 52 recorded neurons, none was expressing ChR2 (checked with a 1s light pulse), and 76.92% of them responded to the light stimulation protocol, meaning they were receiving monosynaptic excitation from aIC or pIC terminals.

### QUANTIFICATION AND STATISTICAL ANALYSIS

Data collection and statistical analysis were performed using Matlab, Microsoft Excel, and GraphPad Prism 6 software. We used one-way ANOVA test and Tukey’s post-hoc tests, two-tailed Student’s *t*-test, and chi-square test. P values of ≤0.05 were considered statistically significant at a confidence interval of 95%. We used Fiji (Image J, NIH) to analyze fluorescent area and cell number. For measurements of CB_1_-fluorescent area in the amygdala, we first change the image to 8-bit and adjust the threshold with a range from 40 to 80. Then, we selected CeA or BLA and measured the fraction of CB_1_-fluorescent area. For counting number of CB_1_-positive cells or IC-BLA projection cells, we selected the IC area and counted cells by using Maxma. Prominences are higher than 40 and 80 for quantification of Low and High CB_1_ cells, respectively. For quantification of colocalization of CB_1_ positive cells and IC-BLA projection neurons, we identified the colocalization by eyes and used Point Tool in Fiji to count cells. For detailed statistical analysis, see statistical tables **(Table S1)**.

## SUPPLEMENTAL INFORMATION

**Table S1**. **Statistical details relate to Figures 1-4, S1, and S3-S4.**

**Video S1. Calcium levels in aIC-BLA projection neurons decrease in response to water licking, related to Figure 3.**

**Video S2. Calcium levels in pIC-BLA projection neurons increases in response to water licking, related to Figure 3.** Water licking is indicated by the blinking LED on the bottom left of the box.

## ACKNOWLEDGMENTS

We thank the animal facility and the genotyping platform of the NeuroCentre Magendie (INSERM U1215 Unit) for assisting in animal breeding, maintenance, and genotyping. We also thank Dr. Anes Ju for helping in analyzing electrophysiological data. The microscopy was done in the Bordeaux Imaging Center a service unit of the CNRS-INSERM and Bordeaux University, member of the national infrastructure France BioImaging supported by the French National Research Agency (ANR-10-INBS-04), which provided the confocal microscope (Leica TCS SP8) and the slide scanner (Nanozoomer 2.0HT, Hamamatsu Photonics France), the help of Sébastien Marais is acknowledged. HHMI Janelia farm research campus is acknowledged for providing the rAAV2-retro helper. We thank the Viral Vector Facility (VVF) of Neuroscience Center Zurich (ZNZ) for providing the rAAV2-retro viral vectors. This work is supported by the China Scholarship Council (to Z.Z.), NARSAD postdoctoral fellowship (to A.C.), INSERM (to G.M., A.B., and L.B.), Nouvelle Aquitaine Region (to G.M. and A.B), European Research Council (Endofood, ERC–2010–StG–260515 and CannaPreg, ERC-2014-PoC-640923, MiCaBra, ERC-2017-AdG-786467, to G.M.), Fondation pour la Recherche Medicale (FRM, DRM20101220445, to G.M.), the Human Frontiers Science Program, the ‘Agence Nationale de la Recherche’ (ANR, NeuroNutriSens ANR-13-BSV4-0006, ORUPS ANR-16-CE37-0010-01, CaCoVi ANR-18-CE16-0001-02, to G.M. and mitoCB_1_-fat-19-JCJC to L.B.).

## AUTHOR CONTRIBUTIONS

Z.Z. and A.B. conceived the project. Z.Z., A.Covelo, L.B., G.M., and A.B. designed the experiments and analyzed data. Z.Z. performed the behavioral and imaging experiments. A.B. and A.Covelo performed *ex vivo* recording experiments. M.V. performed fluorescent *in situ* hybridization experiments. A.M., Y.W., and D.J. performed fiber photometry experiments and analyzed data. A.Cannich prepared reagents. G.M. and A.B. supervised the project. Z.Z., G.M., and A.B. wrote the manuscript. All authors read and approved the manuscript.

